# Chronic lithium treatment alters the excitatory/inhibitory balance of synaptic networks and reduces mGluR5-PKC signaling

**DOI:** 10.1101/2020.09.18.303578

**Authors:** A. Khayachi, A. R. Ase, C. Liao, A. Kamesh, N. Kuhlmann, L. Schorova, B. Chaumette, P. Dion, M. Alda, P. Séguéla, G.A. Rouleau, A. J. Milnerwood

**Author notes:** **Corresponding Authors:** Dr Austen J. Milnerwood, PhD & Dr. Guy A. Rouleau, MD, PhD, FRCPC, OQ, Department of Neurology and Neurosurgery, McGill University, Montréal, Québec, Canada H3A 2B4, &.

## Abstract

Bipolar disorder (BD) is characterized by cyclical alternations between mania and depression, often comorbid with psychosis, and suicide. The mood stabilizer lithium, compared to other medications, is the most efficient treatment for prevention of manic and depressive episodes. The pathophysiology of BD, and lithium’s mode of action, are yet to be fully understood. Evidence suggests a change in the balance of excitatory/inhibitory activity, favouring excitation in BD. Here, we sought to establish a holistic appreciation of the neuronal consequences of lithium exposure in mouse cortical neurons and identify underlying mechanisms. We found that chronic (but not acute) lithium treatment significantly reduced intracellular calcium flux, specifically through the activation of the metabotropic glutamatergic receptor mGluR5. This was associated with altered phosphorylation of PKC and GSK3 kinases, reduced neuronal excitability, and several alterations to synapse function. Consequently, lithium treatment shifts the excitatory/inhibitory balance in the network toward inhibition. Together, the results revealed how lithium dampens neuronal excitability and glutamatergic network activity, which are predicted to be overactive in the manic phase of BD. Our working model of lithium action enables the development of targeted strategies to restore the balance of overactive networks, mimicking the therapeutic benefits of lithium, but with reduced toxicity.

## INTRODUCTION

Bipolar disorder (BD) is a major psychiatric illness affecting 1-3% of the population worldwide and one of the top 10 causes of disability (1, 2). BD starts in adolescence and has a life-long course, characterized by frequently disabling episodes of mania and depression, often associated with psychosis and suicide (3). The etiology of BD is complex and unclear, with genetic and environmental factors implicated. Currently, except for very rare cases, no monogenic cause has been consistently identified, and genome-wide association study (GWAS) hits have been found in varied biological processes, including pathways related to intracellular signal transduction, glutamate synaptic function, hormone signaling, and immune system regulation (4–6). The unknown cause and apparent genetic heterogeneity of BD are a challenge to research efforts, especially those aimed at developing appropriate disease models.

Lithium is an effective treatment for mania and has been consistently shown to reduce suicide and overall mortality (7). Despite its narrow therapeutic range and potential side effects such as tremor, polyuria, decreased thyroid function, and renal toxicity in a minority of patients (8), lithium remains the first line treatment for prevention of both manic and depressive episodes in BD. For many patients, it is the most effective mood stabilizer (9, 10), and in addition to reducing suicide risk, it often enables patients to regain social and occupational function (7). Despite the widespread use of lithium as a BD treatment for over 60 years, the mode of action needs to be better understood, as does the reason why it is effective in only ~30% of BD patients (11, 12).

Studies aimed at elucidating lithium’s mode of action have found macroscopic changes in brain structure (13) and alterations at the cellular level (14). For the latter, it has been shown that acute lithium administration increases glutamate signaling (15, 16). In contrast, longer-term chronic treatment over 6-7 days confers protection against glutamate-induced excitotoxicity, by reducing NMDA receptor-dependent calcium flux (17). It has also been shown that lithium alters various intracellular signaling cascades, by decreasing second messenger and calcium signaling, while inhibiting several enzymes and kinases such as glycogen synthesis kinase 3 (GSK3), extracellular-regulated kinase (ERK)/mitogen-activated protein kinase (MAPK) and inositol monophosphate phosphatase (IMPase) (14, 18, 19). The resulting alterations to intracellular signaling likely also converge upon the regulation of gene expression, synaptic transmission and plasticity, neuroprotection, and circadian biology.

Several genomic studies correlated specific loci with lithium responsiveness (20), suggesting a shared genetic predisposition both to disease and response to treatment. Single nucleotide polymorphisms (SNPs) in the *PLCG1* gene have been associated with response to lithium, indicating that the phospholipase C (PLC)-phosphatidylinositol4,5-biphosphate (PIP2)-inositol triphosphate (IP3) signaling pathway may be an important target of lithium. Interestingly, our unpublished data (72) report an association between a SNP in the *GRM5* gene encoding the metabotropic glutamate receptor 5 (mGluR5) and response to lithium suggesting that mGluR5 activity and downstream PLC-IP3 signaling is also involved in lithium’s therapeutic action. Other studies have found SNPs associated with BD in the *GRIN2A(6)* and *GRIA2* genes that encode NMDA and AMPA receptor subunits, respectively. Intriguingly, only SNPs in GRIA2 were associated with lithium responsiveness (5, 21), suggesting that lithium alters the regulation of Ca^2+^-permeable AMPA receptors. Beyond genomic association, it remains unclear how lithium interacts with mGluR5, PLCG1 and GluA2 signaling to produce the beneficial outcome in lithium-responsive patients.

Here, we sought to establish a holistic appreciation of the neuronal consequences of chronic lithium exposure in mouse cortical neurons and begin to determine the underlying mechanisms. We performed messenger RNA (mRNA) sequencing in neurons treated chronically with lithium and discovered altered transcriptional regulation of genes involved in glutamate receptor trafficking and intracellular calcium signaling. We found that chronic (but not acute) lithium treatment significantly reduced excitatory receptor-mediated intracellular calcium flux, specifically through the mGluR5 receptor. This was associated with altered phosphorylation of PKC and GSK3 kinases, reduced neuronal excitability, and several alterations to synapse function. Specifically, chronic lithium exposure reduced excitatory synapse activity and density, while increasing inhibitory synapse activity and density. Consequently, lithium treatment altered the excitatory/inhibitory (E/I) balance in the network, favouring inhibition. Together, the results shed light on how lithium may dampen neuronal excitability and glutamatergic network activity, which are predicted to be overactive in the manic phase of BD (22–24).

In addition, this discovery strengthens the potential clinical use of lithium to treat disorder with altered excitatory/inhibitory network activity such as epilepsy and several forms of autism. This study could also help to develop targeted strategies to restore the balance of overactive networks, mimicking the therapeutic benefits of lithium, but with reduced toxicity.

## MATERIALS AND METHODS

### Primary neuronal cultures and animals

Cortical neurons were prepared from wild-type (WT) embryonic (E15.5) C57BL/6 mice as previously described (25). Animals were maintained within the Centre for Neurological Disease Modeling according to the Canadian Council on Animal Care regulations (AUP 2017-7888B). Briefly, cortical neurons were plated in Neurobasal medium (ThermoFisher 21103049) supplemented with 1x B27 (ThermoFisher 1750044), 1x glutaMax (ThermoFisher 35050061) on 60-mm dishes or 12-mm glass coverslips (VWR) pre-coated with poly-D-Lysine (0.1 mg mL-1; Sigma). Neurons (600,000 cells per 60-mm dish or 80,000 cells per 12-mm coverslip) were then used at 18-20 days *in* vitro (DIV).

### Drug treatment

The therapeutic range of LiCl (lithium) treatment is between 0.75-1.5mM. In this study, neurons were treated chronically with ~1.5mM LiCl (Sigma L9650) for 7 days starting at 11 DIV post-differentiation. As controls, neurons were treated with ~1.5mM of NaCl (Sigma S5886) to keep the same amount of chloride in the dish as neurons treated with LiCl. Experiments were performed at 18 to 20 DIV.

### Data manipulation and statistical analyses

Statistical analyses were performed using GraphPad Prism software (GraphPad software, Inc). All data are expressed as mean ± standard error of the mean (s.e.m.). Paired t-tests (Fig 5A,B,C), parametric unpaired t-tests (Figs: 2C-G; 5D-F; Supplementary Figs: 3B; 4C,f; 5A-C) or non-parametric Mann-Whitney tests (Figs: 2H; 3B-E; 4; Supplementary Fig. 2A-C) were used to compare medians of two sets. One-sample t-tests were used with hypothetical value 100 for control (Figs: 3G; 5G, H; Supplementary Fig: 2D, E). Normality for all groups was verified using the Shapiro-Wilk tests and p<0.05 was considered significant.

### Data availability

All relevant data are in the figures and supplementary figures. Raw data could be requested from the corresponding author.

A detailed “Materials and Methods” section can be found in the supplementary information.

## RESULTS

### Chronic lithium treatment alters the expression of genes involved in synaptic activity, calcium signaling and neuronal excitability

To identify the cellular processes altered by long-term exposure to lithium, we used a concentration designed to match those used in clinical practice. We examined whether there were detectable changes to neuronal gene expression induced by chronic lithium (cLiCl) treatment, using whole transcriptome sequencing in cortical neuron cultures at 18 DIV. Thirty genes were differentially expressed following 7 days of cLiCl treatment, relative to control (Fig. 1a and Supplementary Fig. 1). Using Reactome and GO analyses of gene clusters, we determined which pathways were significantly altered by cLiCl treatment. We identified pathways associated with trafficking of AMPA receptors, glutamate binding, activation of AMPA receptors and synaptic activity (P=9.7E-5). Calcium ion signaling (P=4.9E-4), CREB phosphorylation and RAS signaling (P=2.5E-4 and P=2.8 E-4) pathways were also implicated, as were pathways related to neurotransmission by chemical synapses (P=1.1E-3; Fig. 1B).

**Figure 1:**
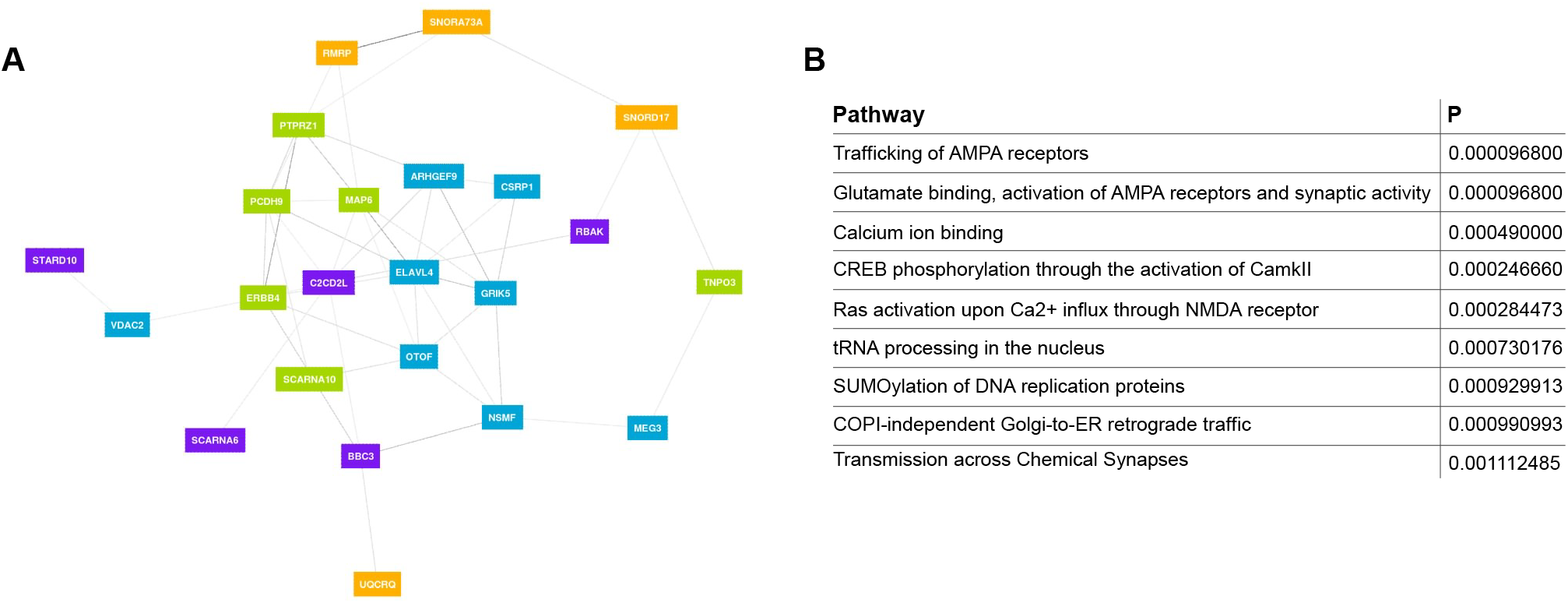
Chronic lithium treatment alters gene expression in mouse cortical neurons. **A)** Gene clustering of differentially expressed genes of primary cortical mouse neurons at 18 DIV treated chronically with LiCl (1.5mM) for 7 days compared to controls. Each gene cluster is identified by a color. **B)** Significant pathways of gene clusters identified through gene network analysis for lithium treatment effect on primary cortical mouse neurons.

### Sub-toxic lithium treatment reduces spine density and alters dendritic spine morphology

The therapeutic window for lithium is very narrow (0.5 to ~1.5mM) and the line between efficacy and toxicity is fine. It is proposed that lithium is toxic above 2mM (26); thus, we confirmed that our chronic lithium treatment at ~1.5mM for 7 days had no detectable toxicity, as assayed by a cell viability test, in primary cortical neurons (Fig. 2C).

**Figure 2:**
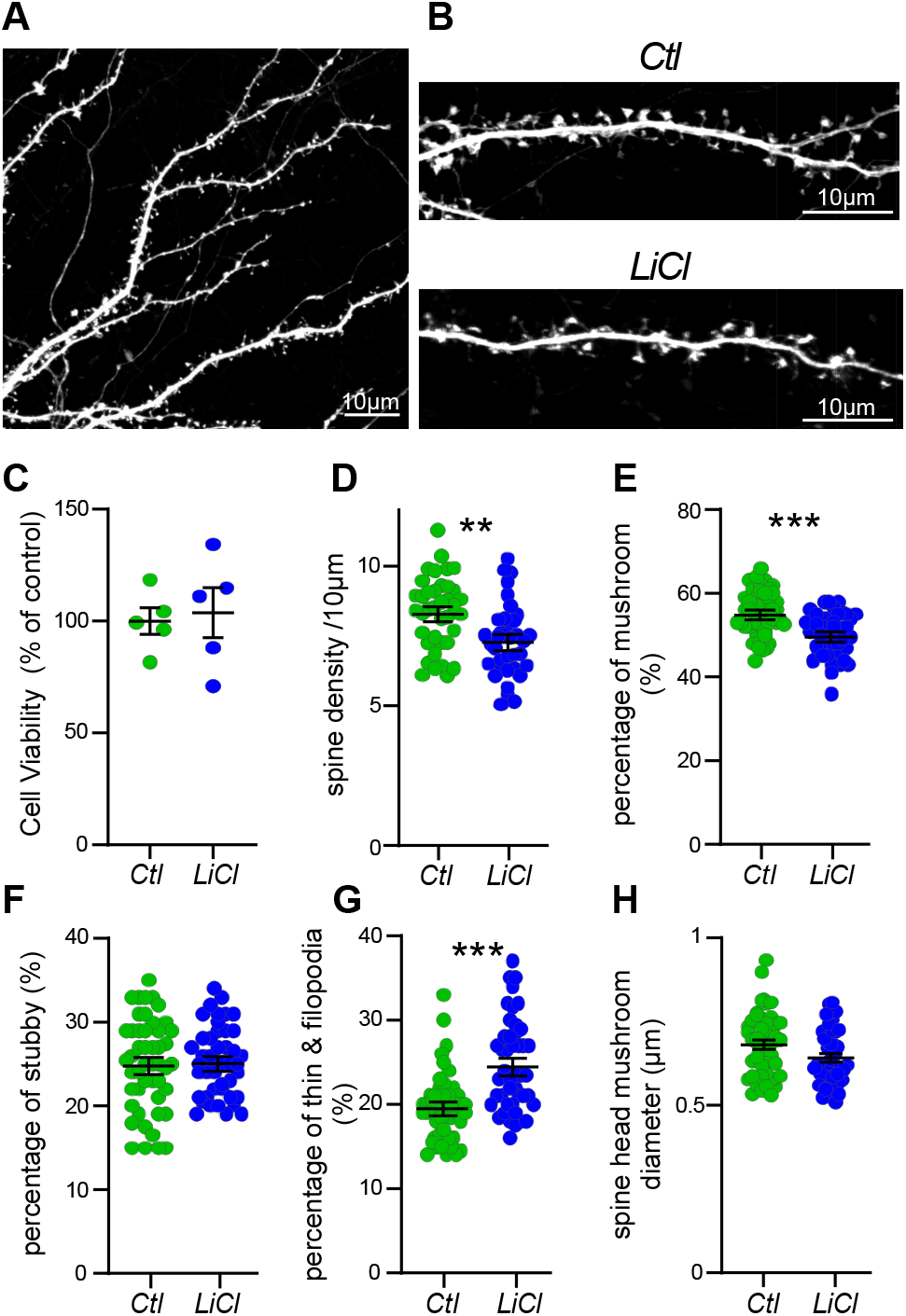
Chronic lithium treatment leads to changes in spine morphology. **A)** Representative confocal image of dendrites expressing free GFP, bar 10μm. **B)** enlargement of a dendrite expressing free GFP treated or not with LiCl (1.5mM) for 7 days, bar 10μm. **C)** Cell viability test shows no toxic effect of LiCl (1.5mM) on neurons treated for 7 days, from 5 independent experiments. Scatter plots show the density of protrusions **(D)** and the relative proportion of mushroom spines **(E)** stubby spines **(F)** and thin spines **(G)** treated or not with LiCl for 7 days. **H)** Scatter plots show the spine head mushroom diameter from neurons treated or not by LiCl. Data shown in **C-H** are the mean ± s.e.m. and statistical significance determined by parametric unpaired t-test for **C-G** and non-parametric Mann-Whitney test for **H**. N=~5000 protrusions per condition from ~45 neurons from four independent experiments, \*\*\**p* < 0.0005, \*\**p* = 0.0013.

Excitatory synapses develop their specialized synaptic structures as they mature, over a similar timeframe *in vivo* and in primary culture. Immature postsynaptic protrusions, filopodia and thin spines, re-appear on dendrites between 4 & 7 DIV after excitatory neurites have regenerated, and new contacts begin to form between axons and dendrites. As postsynaptic structures mature, they become shorter, fatter, and mushroom-like. By 21 DIV spine densities stabilize, with 80-90% of protrusions exhibiting mature morphology (27–29). Several studies have suggested lithium treatment leads to dendritic spines morphological changes (30–32). To determine whether chronic lithium treatment affects dendritic spine density and maturation, we analyzed the presence and morphology of dendritic protrusions in GFP-filled neurons after cLiCl treatment (Fig. 2A, B).

cLiCl treatment slightly reduced the density of protrusions in mouse cortical neurons (Fig. 2D; control 8.34 ±0.21; cLiCl 7.33 ±0.21 / 10μm) suggesting a reduction in the number of excitatory synapses. Analyses of the morphology of the remaining protrusions showed a significant reduction of ~5.3% in the number of mature spines (mushroom spine type, see supplementary Methods for spine characterization guidelines; Fig. 2E). No changes occurred in the number of stubby spines (~25%), but the percentage of immature spines (thin and filopodia spines) increased by ~5% in cLiCl-treated neurons (Fig. 2F, G), matching the reduction in mature protrusions. In addition, we observed a tendency towards a reduction in the mean head diameter of mature spines with cLiCl treatment, from ~0.68μm to ~0.64μm (Fig. 2H). Together, the data show that cLiCl treatment results in smaller and fewer mature spines, indicating that lithium affects either spine maturation or maintenance of spine maturity in primary mouse cortical neurons.

### Lithium induces excitatory and inhibitory synaptic changes

To examine whether the cLiCl induced changes to synapse densities reflecting the results obtained with spines in Fig. 2, we assayed for pre- and post-synaptic markers of excitatory and inhibitory synapses. To estimate excitatory synapse number, we quantified the density, intensity and colocalization of the presynaptic vesicular glutamate transporter 1 (VGluT1) and postsynaptic density protein 95 (PSD95; Fig. 3A, C). PSD95 puncta density was significantly reduced by cLiCl treatment (control 18.9 ± 1.5; cLiCl 12.9 ± 0.9 puncta/10μm), in agreement with the reduction in protrusion density and number of mature spines in neurons treated with cLiCl (Fig. 3C, D). Further, there were significantly fewer VGluT1/PSD95 co-clusters in cLiCl-treated neurons (control 5.9 ± 0.5; cLiCl 4.5 ± 0.4 co-clusters/10μm; Fig. 3D), indicative of a reduction in excitatory synapse number after cLiCl treatment. VGluT1 (and VGAT, see below) puncta intensity was increased in remaining cLiCl-treated clusters (Supplementary Fig. 2B, C; VGlut1: control: 10.97 ± 0.3; cLiCl: 13.41 ± 0.5 a.u; VGAT: control, 12.91 ± 0.4; LiCl, 17.07 ± 0.6 a.u). Conversely, PSD95 puncta intensity was reduced in cLiCl-treated cultures (control, 6.99 ± 0.4; cLiCl: 5.67 ± 0.2 a.u.; Fig. 3E), in agreement with the reduction of the mean head diameter of mature spines seen in Fig 2H. Puncta density and colocalization of the inhibitory presynaptic vesicular GABA transporter (VGAT) and postsynaptic GABA receptor scaffold Gephyrin were also quantified (Fig. 3A, B). We observed a significant increase in the density of Gephyrin puncta, and VGAT/Gephyrin co-clusters, in cLiCl-treated neurons (control: 7.34 ± 0.5; cLiCl: 11.56 ± 1.4 Gephyrin puncta/10μm; control: 1.87 ± 0.09; cLiCl: 2.58 ± 0.2 of VGAT/Gephyrin co-clusters/10μm; Fig. 3B). Gephyrin puncta intensity was unchanged but VGAT puncta density was increased in cLiCl-treated neurons (supplementary Fig. 2A, B). The data indicate that lithium treatment increased the number of inhibitory synapses.

**Figure 3:**
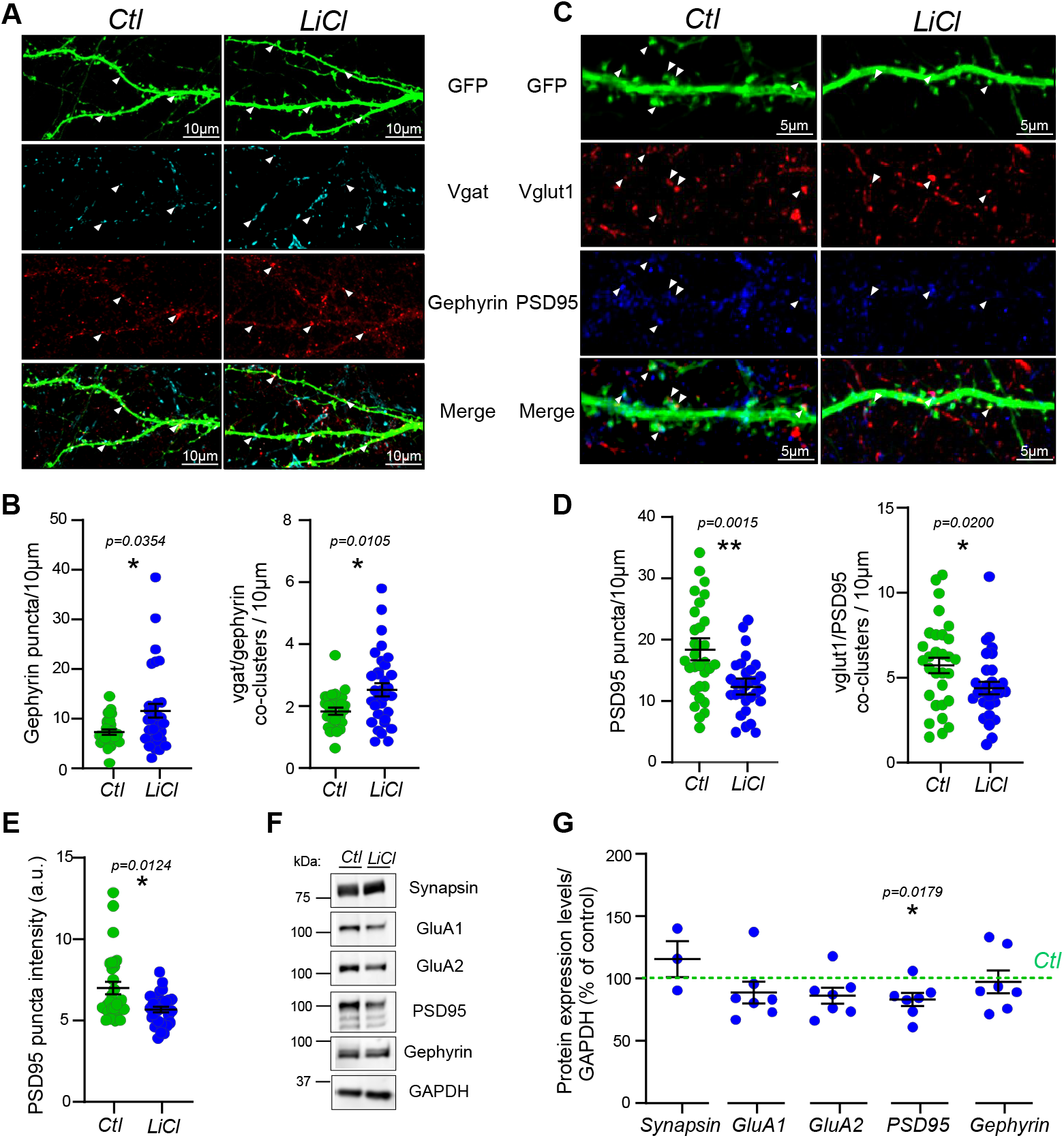
Chronic lithium treatment induces excitatory and inhibitory synaptic changes. **A)** Representative confocal image of dendrites expressing free GFP from neurons treated or not with LiCl (1.5mM) for 7 days, with antibodies directed against Vgat and Gephyrin. Arrowheads show the Vgat and Gephyrin puncta localization and the co-localization between Vgat and Gephyrin in the merge indicating the inhibitory synapses. Bar, 10μm. **B)** Scatter plots show quantification of Gephyrin puncta density/10 μm and Vgat-Gephyrin co-cluster density representing the density of inhibitory synapses per 10μm from secondary/tertiary dendrites from neurons treated or not with LiCl (1.5mM) for 7 days. N = 32 neurons per condition from three separate experiments. **C)** Representative confocal image of dendrites expressing free GFP from neurons treated or not with LiCl (1.5mM) for 7 days, with antibodies directed against Vglut1 and PSD95. Arrowheads show the Vglut1 and PSD95 puncta localization and the co-localization between Vglut1 and PSD95 in the merge indicating the excitatory synapses. Bar, 5μm. **D)** Scatter plots show quantification of PSD95 puncta density/10 μm and Vglut1-PSD95 co-cluster density representing the density of excitatory synapses per 10μm from secondary/tertiary dendrites as well as PSD95 puncta intensity **E** from neurons treated or not with LiCl (1.5mM) for 7 days. N = 30 neurons per condition from three separate experiments. Data shown in **B-E** are the mean ± s.e.m. and statistical significance was determined using a non-parametric Mann-Whitney test. **F)** Representative immunoblots anti-Synapsin1, GluA1, GluA2, PSD95, Gephyrin and GAPDH of 18 DIV neuronal extract from neurons treated chronically or not with LiCl (1.5mM). **G)** Quantification with scatter plot of some presynaptic and postsynaptic protein expression levels normalized with GAPDH and represented as percentage of control of 18 DIV neuronal extract from neurons treated chronically or not with LiCl (1.5mM) from three to seven separate experiments. Data show the mean ± s.e.m and statistical significance was determined using one sample t test with hypothetical value 100 for control.

To further assess the reduction in number of excitatory synapses (Fig. 3D) induced by cLiCl, and determine whether other synaptic changes are occurring, we measured protein levels of PSD95, GluA1 and GluA2 AMPAR receptor subunits by western blot (Fig. 3F, G). GluA1 protein was unchanged, whereas GluA2 and PSD95 were significantly reduced in response to cLiCl (−13.8% GluA2 and −16.9% PSD95 of controls; Fig. 3G). This data suggests that cLiCl treatment downregulates GluA2-containing AMPA receptors, as well as the number of excitatory synapses. Interestingly, the results were accentuated by a higher dose of cLiCl (3.5mM; −54.42% GluA2 and −54.49% PSD95 of controls; Supplementary Fig. 2D). Notably, the expression level of AMPA receptors and PSD95 were unchanged when neurons were treated acutely (aLiCl,1.5mM) for 4h (Supplementary Fig. 2E), suggesting longer time is needed for LiCl to alter the levels of these proteins. The expression levels of Synapsin1 and Gephyrin were unchanged by cLiCl or aLiCl (1.5mM, Fig. 3G and Supplementary Fig. 2D, E).

### Lithium decreases neuronal excitability and excitatory synaptic transmission, while increasing inhibitory synaptic transmission

To examine the functional consequence of lithium treatment on neuronal networks, we assessed intrinsic membrane excitability and action potential generation, in addition to quantification of excitatory and inhibitory synaptic transmission. To assess neuronal excitability, we recorded membrane deflection in response to current injection (Supplementary Fig. 3B) and action potential (AP) firing induced by depolarizing currents in current clamp. Although highly variable, cLiCl-treated neurons appeared to fire fewer APs than control neurons (Fig. 4A-C and supplementary Fig. 3A), indicating that cLiCl reduces cell excitability. We then assayed sodium and potassium currents in voltage clamp and found both were reduced in cLiCl-treated neurons. Specifically, peak sodium current was −16.3% of control (Fig. 4D, E; control: 4.97± 0.3 pA; cLiCl: 4.16± 0.3 pA), and slow and fast potassium currents were reduced −19.3% and −12.13% in cLiCl-treated neurons, compared to controls (Slow K current; control: 2.6± 0.15 pA; cLiCl: 2.1± 0.13 pA; Fast K current; control: 3.3± 0.19 pA; cLiCl: 2.9± 0.16 pA; Fig. 4F-H, and supplementary Fig. 3C) These data demonstrate that cLiCl treatment alters sodium and potassium channel conductances, and decreases membrane excitability.

**Figure 4:**
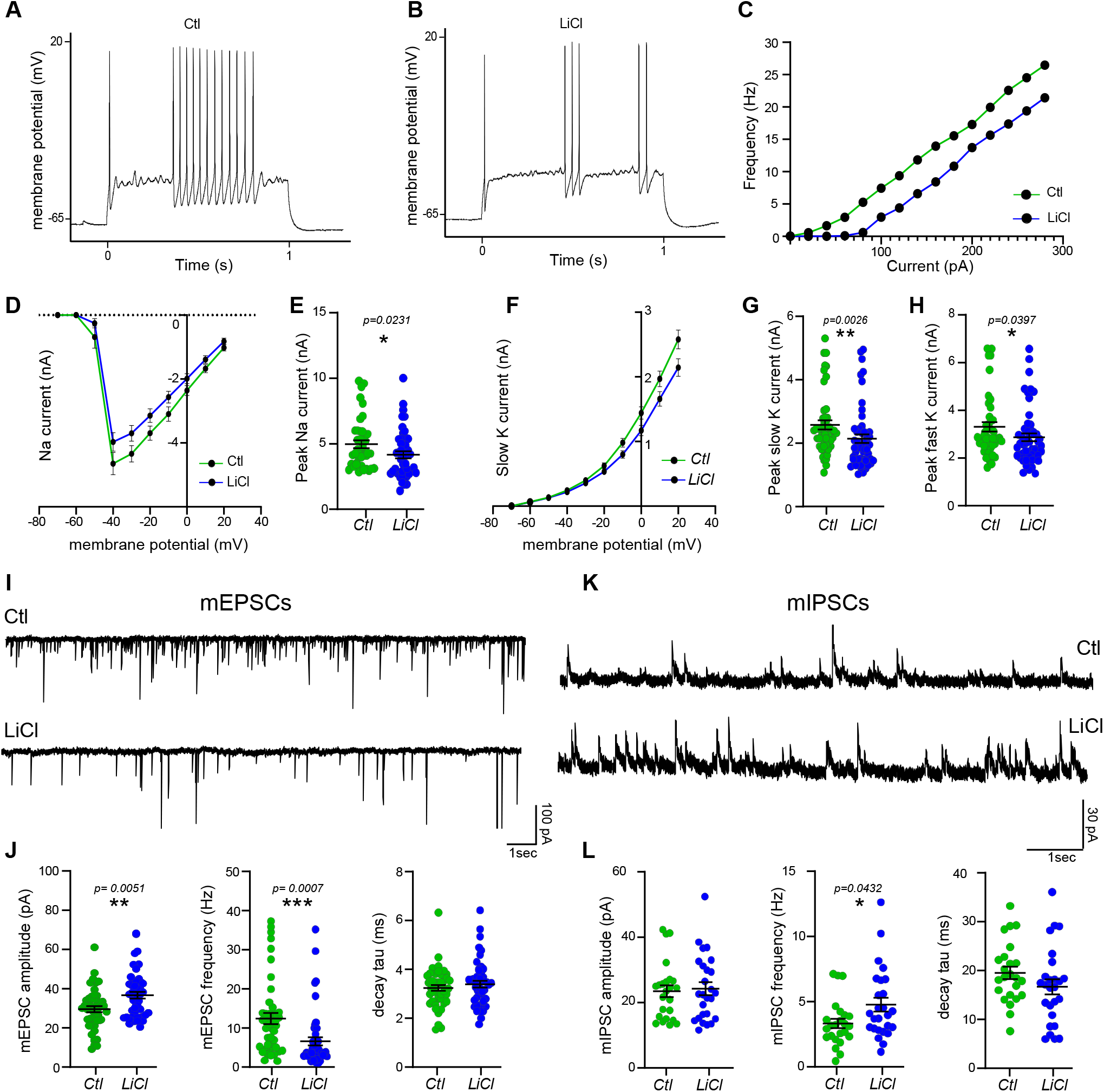
Chronic lithium treatment reduces neuronal excitability and excitatory transmission, while increasing inhibitory synaptic transmission. Representative action potential trains in control **(A)** and chronically LiCl (1.5mM) treated neurons **(B)** at DIV 18, in response to a 1 second depolarizing 120 pA current step from ~-65mV. **C)** Frequency-current (F-I) plot among repetitively-firing neurons. Frequency represents the mean number of spikes/second from ~32 neurons per condition from four independent experiments. Voltage dependence of the amplitude of the sodium current in **D** and quantification of the peak amplitude of sodium currents in neurons treated or not chronically with LiCl (1.5mM) in **E**. Voltage dependence of the amplitude of the slow potassium current in **F** and quantifications of the peak amplitude of the slow and fast potassium currents in neurons treated or not chronically with LiCl (1.5mM) in **G** and **H**. The data from **D-H** are from five separate experiments and show the mean ± s.e.m and statistical significance was determined by a non-parametric Mann-Whitney test. **I)** Representative sample traces of mEPSCs from neurons treated or not with LiCl (1.5mM) for 7 days. Scale bar showed as inset. **J)** Scatter plot show quantification of amplitude, frequency and decay tau of mEPSCs of ~45 neurons from four independent experiments. **K)** Representative sample traces of mIPSCs from neurons treated or not with LiCl (1.5mM) for 7 days. Scale bar showed as inset. **L)** Scatter plot show quantification of amplitude, frequency and decay tau of mIPSCs of ~26 neurons from three independent experiments. Data shown in **J** and **L** are the mean ± s.e.m. and statistical significance determined by non-parametric Mann-Whitney test.

Synaptic network activity was assessed by recording quanta of AMPAR-mediated miniature excitatory, and GABAR-mediated miniature inhibitory post-synaptic currents (mEPSCs and mIPSCs) by voltage-clamp recording (Fig. 4I, K). In neurons treated with cLiCl, mEPSC event amplitude was 27% increased compared to control neurons (control: 29.67 ± 1.5pA; cLiCl: 37.71 ± 1.9pA) and event frequency was reduced by 47% (control: 12.44 ± 1.4 Hz; cLiCl: 6.5 ± 1 Hz; Fig. 2J and supplementary Fig. 4A, B), while no changes were observed in event decay tau (Fig. 4J). Event amplitude was not different for mIPSCs (control: 23.46 ± 1.8 pA; cLiCl: 24.25 ± 1.9 pA), but there was a significant increase in mIPSC event frequency in cLiCl treated cultures (control: 3.3 ± 0.35 Hz; cLiCl: 4.8 ± 0.53 Hz; Fig. 4L and supplementary Fig. 4D, E), and again no change in event decay tau (Fig. 4l). Measures of membrane properties of voltage-clamp recordings can be found in supplementary Fig 4C, F. Together, electrophysiological experiments demonstrate that cLiCl treatment alters synaptic network properties by decreasing excitatory and increasing inhibitory activity.

### Chronic lithium treatment downregulates mGluR-mediated calcium response and signaling

Since cLiCl treatment downregulated the major depolarizing AMPAR current at glutamatergic synapses, it is of interest to determine which regulatory signal transduction pathways are altered, and whether any changes in these are a consequence, or cause, of altered synaptic transmission. To this end, we investigated ionotropic and metabotropic glutamate receptor-mediated calcium signaling, in cultures acutely or chronically treated with LiCl. Ratiometric calcium imaging was conducted to measure intracellular calcium levels and flux upon stimulation. Tetrodotoxin (TTX) was added to block sodium channels and action potential burst firing, thus calcium flux was directly in response to glutamate receptor activation. The intracellular calcium level at rest was similar between cLiCl and control-treated neurons (Supplementary Fig. 5A). Control neurons were first exposed to repeated glutamate pulses, which did not attenuate calcium (Ca^2+^) flux upon repeated applications (Supplementary Fig. 5B). Then, following exposure to 1μM glutamate, an acute (5min 1.5mM) LiCl treatment was applied prior to repeated glutamate stimulations. The second calcium response was similar to the first, indicating that LiCl does not directly reduce glutamate-induced Ca^2+^ flux or act as an antagonist of glutamate receptors (Fig. 5A). The use of specific agonists of NMDA (Fig. 5B) and mGluR5 (Fig. 5C) shows that acute LiCl treatment had no differential effect on either NMDA or mGluR5-mediated Ca^2+^ responses (Fig. 5B, C). A medium-term LiCl treatment for 4h similarly did not alter glutamate-induced Ca^2+^ flux (supplementary Fig 5C).

**Figure 5:**
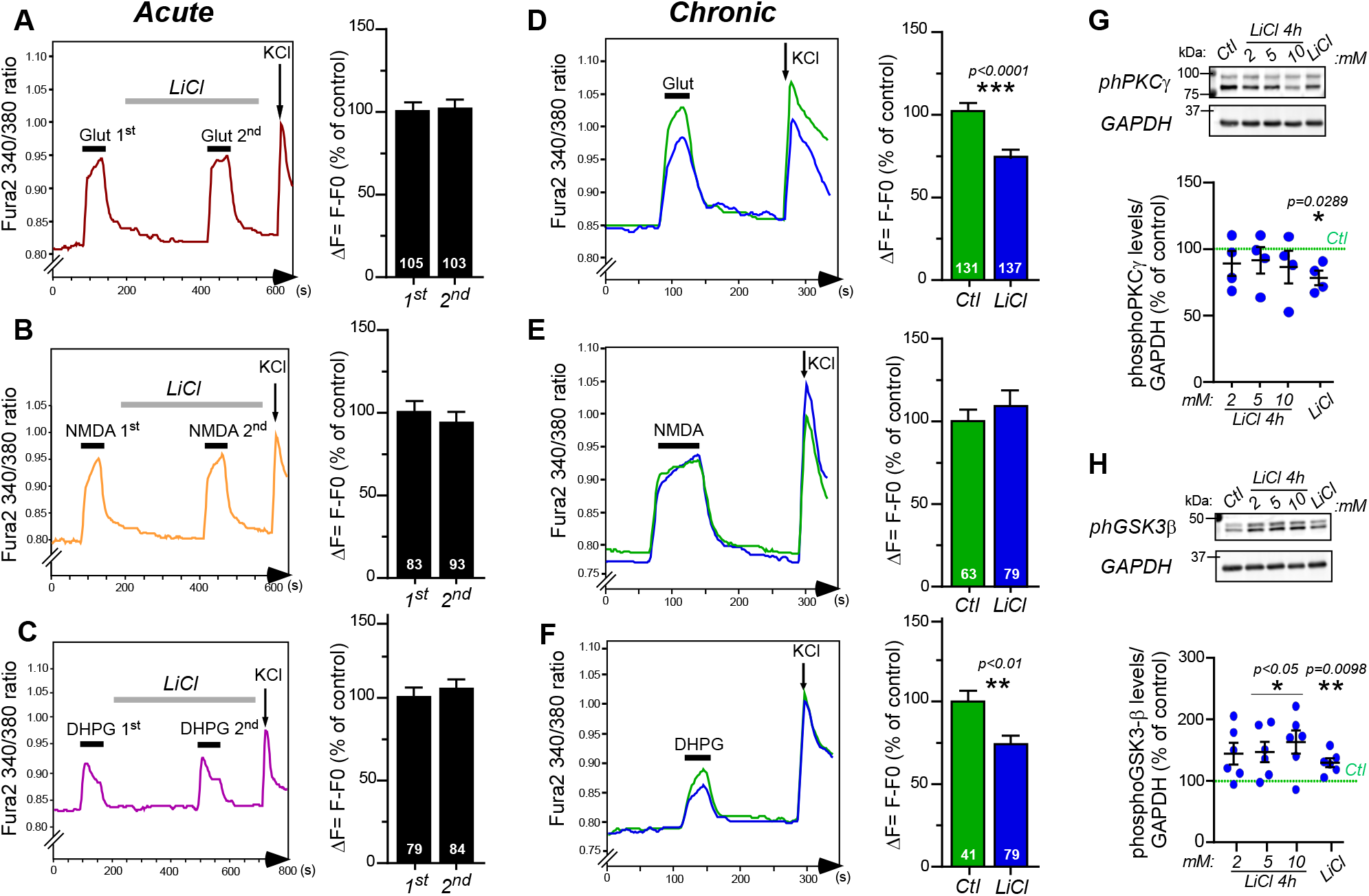
Chronic lithium treatment reduces mGluR-mediated calcium response and signaling. **A-C)** Representative sample traces of calcium responses after glutamate (1μM), NMDA (10μM) and DHPG (100 μM) stimulation with histograms showing the quantification of calcium changes upon stimulation in mouse primary cortical neurons. The second stimulations are preceded by acute LiCl (1.5mM) treatment of 5min, then followed by 2sec of KCl stimulation. **D-F)** Representative sample traces of calcium responses after glutamate (1μM), NMDA (10 μM) and DHPG (100μM) stimulation with histograms showing the quantification of calcium changes as percentage of control upon drugs stimulations in mouse primary cortical neurons chronically treated or not with LiCl (1.5mM) for 7 days. Number of neurons is indicated on each histogram from 3 independent experiments. Data shown in **A-F** are the mean ± s.e.m. and statistical significance determined by paired t-test for acute treatment **(A-C)** and unpaired t-test for chronic treatment **(D-F)**. **G-H)** Representative immunoblots and quantification of anti-phosphorylated levels of PKCγ (Thr514) and GSK-3β (Ser9) normalized with GAPDH and represented as percentage of control from neuronal extract from mouse primary cortical neurons treated with 2, 5 and 10mM of LiCl for 4h, or chronically by LiCl (1.5mM) for 7 days. The data are from four (for phPKC) and six (for phGSK3) separate experiments and show the mean ± s.e.m. and statistical significance determined by one sample t-test with hypothetical value 100 for control.

In contrast to acute application, glutamate stimulation of neurons that were chronically treated with cLiCl (7 days) exhibited a significant decrease in Ca^2+^ response amplitude (cLiCl: ΔF=73.9% of control; Fig. 5D). The amplitude of the Ca^2+^ response upon specific activation of NMDA receptors was unchanged, indicating that chronic LiCl treatment did not affect NMDAR-mediated Ca^2+^ responses (Fig. 5E). Conversely, cLiCl treatment significantly reduced Ca^2+^ response amplitude produced by direct stimulation of mGluR5 (by DHPG agonism; cLiCl, ΔF=70.8% of control; Fig. 5F). The data suggest that cLiCl exposure specifically reduces Ca^2+^ release from the endoplasmic reticulum (ER). This attenuation of Ca^2+^ responses to glutamate and DHPG stimulation could be due to impaired receptor activation e.g., a reduction in the number or sensitivity of mGluR5 receptors at the cell surface, or IP3 receptors on the ER. Alternatively, lithium may affect the mGluR5-PLC-PIP2-DAG-IP3 signaling cascade, i.e. through altered PLC activation, PIP2 hydrolysis or DAG and IP3 availability in the cell. Either possibility would result in reduced ER calcium release.

To determine whether cLiCl-induced attenuation of glutamate-mediated Ca^2+^ responses affects downstream signaling pathways, we assayed a major downstream target of mGluR5 activation, protein kinase C (PKC), which is responsible for regulating a wide range of neuronal function, such as excitability, neurotransmission, and plasticity (33–37). PKCγ isoform is exclusively expressed in the brain, and its activity is regulated by phosphorylation at threonine 514 (phPKCγ). The levels of phPKCγ measured by Western Blot from neurons treated with 2, 5 and 10mM of LiCl for 4h were similar to controls (Fig. 5G). However, phPKCγ levels, were reduced following cLiCl (1.5mM) treatment (a reduction of 21.67%; Fig. 5G). Our data demonstrate that PKCγ activity is not directly affected by LiCl, but is a consequence of prolonged exposure.

A second major downstream effector of mGluR5 stimulation is GSK3β, a kinase involved in several neuronal processes such as cytoskeletal reorganization and neuroplasticity (38). Phosphorylation of GSK3B at the Serine 9 (phGSK3β) residue inhibits GSK3β kinase activity. In contrast to acute LiCl effects on PKC, a 4h LiCl treatment with increasing concentrations (2, 5 and 10mM) and chronic LiCl treatment both significantly increased phGSK3β, in a dose-dependent manner (Fig. 5H). These results demonstrate that LiCl has a rapid, and likely direct, effect on GSK3β kinase activity (LiCl 2mM: 144%; LiCl 5mM: 147%; LiCl 10mM: 163% and cLiCl 1.5mM: 129.5% of controls).

## DISCUSSION

Here, we provide the first holistic report of chronic lithium treatment decreasing the balance of excitatory to inhibitory synaptic transmission in cortical neuron networks. This appeared to be a result of altered intracellular calcium signal transduction, expressed by changes to the number and function of excitatory, and inhibitory synaptic connections.

Throughout development and into adulthood, neural connectivity at synapses is subject to dynamic regulation including formation, maintenance, and elimination of synapses themselves. Synaptic transmission is believed to be the means by which all experiences and motivations are stored and utilized, and synaptic plasticity (rapid, activity-dependent alterations to synaptic transmission) is the leading candidate for the cellular basis of learning and memory. In the mammalian forebrain, most excitatory synapses occur on dendritic spines, and changes to their number, morphology, and activity are modelled by increases (long term potentiation; LTP) and decreases (long-term depression; LTD) in synaptic weighting (39, 40). With LTP, spines enlarge and become more mushroom-like, whereas LTD is associated with spine shrinkage. Disruptions to dendritic spine shape, size or number accompany many neurodegenerative diseases, and it has been suggested that dendritic spine alterations are also the substrate of many neuropsychiatric disorders, particularly those that involve deficits in information processing, such as autism spectrum disorder and schizophrenia (40).

Decreased spine density has been shown *post-mortem* in a preliminary study of the subiculum in mood disorder patients (41), and the prefrontal cortex of BD patients (42). It is unclear how reduced spine density in these individuals might reflect a general state of the condition, a brain region-specific effect, or whether it could even be a result of successful medication. In our hands, lithium reduced measures of excitatory spine maturity, and appeared to utilize processes similar to those employed during physiological LTD, decreasing excitatory activity and the number of mature *vs*. immature spines. The increase of mEPSC amplitude observed here in neurons treated with cLiCl could reflect the increase of immature spines phenotype as suggested in previous studies (43–45).

In support of our results, chronic lithium treatment reduced spine density in a Fragile X mouse model (30). However, spine density was shown to be increased with acute lithium treatment in a DIXDC1 KO mouse model of depression (31). It may be that lithium generally facilitates network rearrangements and normalizes spine dysfunction in whichever direction is required, but further investigations may provide consensus on which direction and over what time-frame changes usually occur. It has also been proposed that lithium rescues spine pathology in BD by reducing phosphorylation of the cytoskeleton regulator collapsing response mediator protein-2 (CRPM2) (32), in rat hippocampal cultures, chronic treatment with a dose 2x higher than here, resulted in enlarged spines and increased spine density. Here, we found that chronic lithium not only reduced spine number, but also decreased the percentage of mature spines, mature spine width, and PSD95 puncta intensity. These discrepancies may be due to the higher dose and/or different responses between rat hippocampus and mouse cortex. More studies will be required to settle this discrepancy.

Several rodent models of mania have been generated which traditionally relied upon pharmacological (e.g., psychostimulant amphetamine-induced) or environmental (sleep deprivation-induced) stresses to induce mania-like states, and more recently several transgenic mice have been developed (46, 47). A recent study using knock-in mice of the Ank3 W1989R (48), a variant reported as carried by a BD family (and found in approximately 1:10,000 European Americans) (49), showed a reduction in mIPSCs and an increase in mEPSC frequency, leading to neuronal hyperexcitability. If this is the case in untreated lithium-responsive patients, then our finding that chronic lithium treatment has the opposite effect may explain its therapeutic effect. This could also explain the efficacy of lithium in BD Ank3 mutation carriers(48). Elsewhere, Yang el al (50) generated forebrain-specific PLCγ1 knock-out mice that exhibited manic-like behavior and cognitive deficits associated with a significant reduction in mIPSC frequency. Here, we show that chronic lithium treatment promotes inhibitory transmission, and increases gephyrin clusters at inhibitory synapses. Together, these studies suggest that manic phases correspond to an increased E/I synaptic ratio. Our results are in line with other studies (51, 52) indicating that chronic lithium treatment can counteract abnormalities in E/I circuit balance observed in the mania state of BD animal models, by rearranging the number, morphology and function of excitatory and inhibitory synapses, in a manner that favours inhibition. On this note, elsewhere, lithium has also been shown to reduce synaptic AMPA receptor expression (53–55), again consistent with the reduction in excitatory synapse number observed here.

An important aspect of our results is the demonstration that chronic effects of lithium are distinct from acute effects. Specifically, lithium treatment from 5min to 4h did not affect glutamate-mediated Ca^2+^ responses, demonstrating that lithium did not act as an antagonist of glutamate receptor transmission in our hands. Conversely, mGluR5 glutamate receptor-mediated Ca^2+^ signaling was specifically reduced in neurons chronically treated with LiCl, suggesting that time is required for lithium to attenuate the mGluR5-PIP2-IP3 pathway. Sourial-Bassilious et al (56) concluded lithium attenuates intracellular Ca^2+^ levels due to the downregulation of mGluR5 expression at the plasma membrane, as well as a decrease in intracellular Ca^2+^ in the ER. Others have suggested lithium inhibits inositol monophosphatase and inositol polyphosphate-1-phosphatase, in addition to the inositol transporter (57). This would reduce PIP2 and IP3 availability in the cell to trigger Ca^2+^ release from the ER. Either way, we sought to find out whether a decrease in Ca^2+^ signaling caused by chronic lithium treatment altered downstream effectors.

Our results demonstrate that chronic and acute lithium inhibit GSK3β kinase activity, in a dose dependent manner, but only chronic treatment reduces PKCγ kinase activity. These two kinases are major regulators of synaptic receptor traffic and function (58), actin cytoskeleton reorganization, neuronal transmission and plasticity, as well as gene expression (37, 38, 59–61). This may be the primary mechanism by which lithium acts to prevent mania in BD, where the glutamatergic system is predicted to be overactivated (22–24) and the inhibitory system downregulated (62, 63). On this background, increased PKC activity and levels have been found *post-mortem* in the frontal cortex of bipolar patients. Furthermore, PKC hyperactivity has been detected in the blood of BD patients (64) and animal models of mania, in agreement with our conclusions here and in support of the potential for PKC inhibitors as therapeutics for mania in BD (59, 65–67). It has been previously shown that lithium can reduce glutamatergic neurotransmission by slowing synaptic vesicle (SV) exocytosis (68). This may explain increased VGlut1 cluster intensity we saw here (Supplementary Fig. 2b, c). Although evidence is lacking for how lithium might alter the kinetics of the SV cycle, we would expect reduced GSK3 and PKC activity to impact presynaptic function. It may also be that intracellular Ca^2+^ signaling, via presynaptic mGluR5 autoreceptors (69) is reduced, which may impair vesicular release (70).

Recent advances in human induced pluripotent stem cell (iPSC) technology have provided the means by which to study an individual patient’s neurons. Recently, Mertens et al (71) discovered hyperexcitability in iPSC-derived hippocampal-like neurons from BD patients, which was reversed by lithium treatment (in cells from BD lithium responders). These tools will facilitate our future studies of lithium’s mode of action, at the cellular and network level, in neurons from BD lithium responders *vs* non-responders. We will also determine whether the working model we provide here operates in human scenarios. Ongoing efforts will examine the mGluR5-PLC-IP3 intracellular Ca^2+^ signaling pathway in neurons from BD lithium responders, and the effects of lithium treatment. We hope this will facilitate development of novel disease-modifying therapies for BD, with the same therapeutic benefit of lithium, but with less side-effects.

## ACKNOWLEDGMENTS

We gratefully acknowledge the financial supports from Fonds de recherche en santé du Québec (FRSQ), Ellen foundation and Killam to A.J.M, the Canadian Institutes of Health Research (CIHR) grant (#332971 to G.A.R), ERA PerMed grant to MA and G.A.R, the Healthy Brains for Healthy Lives (HBHL) and Bettencourt-Schueller fondation grants to BC and the RI-MUHC 2020 fellowship to LS. We also thank the microscopy platform of the Montreal Neurological Institute (MNI) and the Molecular Tools Platform of the CERVO Brain Research Center for providing us the AAV constructs. We would also like to thank the animal care facility of MNI and Bruno Vieira for assistance with mice handling. We also thank Dr Simon Wing lab for the kind gift of reagents. G.A.R. holds a Canada Research Chair in Genetics of the Nervous System and the Wilder Penfield Chair in Neurosciences. C.L. is a recipient of the Vanier Canada Graduate Scholarship from the CIHR.

## AUTHOR CONTRIBUTIONS

AK performed all the spine morphology, density and excitatory/inhibitory synapses analyses. AK, AKam and NK performed the electrophysiological recordings, analyzed by AK. A.R.A performed and analyzed the calcium imaging experiments. CL analyzed the RNA sequencing. LS performed the cell viability test and provided some computational tools to analyze imaging data. AK prepared all neuronal cultures and all biochemical experiments. AK, A.J.M and G.A.R contributed to study design, curation and development, and data interpretation. A.J.M and G.A.R provided the overall supervision and funding. AK wrote the original draft and all authors revised and commented on the manuscript, edited by A.J.M.

## AUTHOR CONTRIBUTIONS

All the authors declare no conflict of interest.

