## supplementary information for "Chronic lithium treatment alters the excitatory/inhibitory balance of synaptic networks and reduces mGluR5-PKC signaling"

*Khayachi et al.,*

**Supplementary Materials and Methods**

**Confocal imaging**

*Image acquisition*: For fixed cells, confocal images (1024 × 1024) were acquired with a ×63 oil immersion lens (numerical aperture NA 1.4) on an SP8 confocal microscope (Leica Microsystems). Z-series of 8 images of randomly selected secondary/tertiary dendrites were compressed in 2D using the max projection function in image j (FIJI).

*Dendritic spine imaging and analysis*: Neurons were transduced at 16 DIV by AAV to express GFP under a human Synapsin promoter for 48h. Cells were then fixed at 18 DIV in 3.7% PFA + 5% sucrose for 1h at RT and mounted in Prolong before confocal examination. About 4200–4800 spines were analyzed per condition (two to four dendrites per neuron and from 35 to 40 neurons per condition from four independent experiments). At the time of acquisition, laser power was adjusted so that all spines were below the saturation threshold. To analyze dendritic protrusions parameters, projection images were imported into NeuronStudio software (1), which allows for the automated detection of mushroom, stubby and thin dendritic spines. The length of individual spines was automatically measured, and data were imported in GraphPad Prism software for statistical analysis. Mushroom spines were characterized by a head diameter ranging from 0.3 to 2 µm and a spine length between 0.3 and 5 µm. Stubby spines were identified by a head diameter ranging from 0.3 to 2 µm and a spine length between 0.1 and 0.8 µm. Finally, thin and filopodia spines corresponded to protrusions with a head diameter below 0.3 µm and a spine length ranging from 0.1 to 6 µm.

*Synapse quantification:* Co-clusters (VGlut1-PSD95 and VGAT-Gephyrin) co-occurring with GFP were quantified with SynapCountJ, an imageJ plugin(2)*.*

*Pre and postsynaptic puncta quantification:* VGlut1, VGAT, PSD95 and Gephyrin puncta co-occurring with GFP were quantified by Cell profiler software(3)*.*

### Immunocytochemistry

Neurons (19–20 DIV) were fixed in phosphate-buffered saline (PBS) containing 3.7% formaldehyde and 5% sucrose for 1 h at room temperature (RT), then in NH4Cl (50mM) for 10 min. Neurons were then permeabilized for 20 min in PBS containing 0.1% Triton X-100 and 10% goat serum (GS) at RT and immunostained with a guinea-pig polyclonal anti-vglut1 (1/4000; Millipore AB5905), a rabbit anti-psd95 clone K28/43 (1/1000; NeuroMab), a rabbit anti-vgat (1/500; Synaptic Systems 131002), a mouse monoclonal anti-gephyrin (1/500; Synaptic Systems 147111), antibodies in PBS containing 0.05% Triton X-100 and 5% GS. Cells were washed three times in PBS and incubated with the appropriate secondary antibodies (1/1000) conjugated to Alexa 594 or Alexa647 and mounted with Prolong (ThermoFisher P36930) until confocal examination.

### Cell viability test

### Cell Counting kit 8 using (2-(2-methoxy-4-nitrophenyl)-3-(4-nitrophenyl)-5-(2,4-disulfophenyl)-2H-tetrazolium, monosodium salt) (WST-8/CCK-8; Sigma 96992) is a cell viability assay. Briefly, the kit uses a water-soluble tetrazolium salt to quantify the number of live cells by producing an orange formazan dye upon bio-reduction in the presence of an electron carrier. WST-8/CCK-8 is added directly to the cells for 1h where it is reduced by cellular dehydrogenases to an orange formazan product that is soluble in tissue culture medium. The amount of formazan produced is directly proportional to the number of living cells and is measured by absorbance at 460 nm.

### Immunoblot

### Neurons at 18 DIV were homogenized in lysis buffer (10 mM Tris–HCl pH7.5, 10 mM EDTA, 150 mM NaCl, 1% Triton X100, 0.1% SDS) in the presence of a mammalian protease inhibitor cocktail (Sigma, 1/100 P8340) and phosphoSTOP (Sigma 4906845001) to protect proteins from dephosphorylation. Protein extracts (20-40μg) were resolved by SDS–PAGE, transferred onto PVDF membrane (Millipore IPFL00010), immunoblotted with the indicated concentration of primary antibodies: rabbit polyclonal anti-psd95 clone K28/43 (1/2000; NeuroMab); rabbit monoclonal anti-GluA1 (1/1000; Millipore 05-855R), rabbit anti-GluA2 (1/2000; Sigma AB1768-I), rabbit anti-SynapsinI (1/1000 Millipore AB1543P), rabbit anti-phospho-GSK3b (1/1000; New England BioLabs 9323), rabbit anti-phospho- PKCγ (Thr514) (1/1000; New England BioLabs 9379). Standard GAPDH loading controls were included using a mouse monoclonal anti-GAPDH antibody (1/2000, ThermoFisher MAB5-15738). Then membrane was revealed using the appropriate LI-COR fluophore-conjugated secondary antibodies. Images were acquired on a LI-COR Odyssey Infrared image system. Fluorescence intensity values for each protein of interest were normalized to GAPDH signal from the same gel. Full-size blots for cropped gels can be found in Supplementary figure 6.

### Electrophysiological recordings and analyses

### All electrophysiological signals were acquired using Multiclamp 700B amplifier digitized at 10kHz and pClamp10 software. Data were analyzed in Clampfit10 (Molecular Devices).

### *External and internal solutions*: Neurons at 18 to 20 DIV on coverslips were transferred to a recording chamber in a recording buffer containing (in mM): 167 NaCl, 10 D-glucose, 10 HEPES, 2.4 KCl, 1 MgCl_2_, and 2 CaCl_2_ (300-310 mOsm, pH adjusted to 7.4 with NaOH). Whole cell patch clamp experiments were carried out at RT (22–25 °C) on pyramidal cell looking from cultured mouse cortical neurons.

### For mEPSC and mIPSC recordings, pipettes were filled with a Cesium based solution containing (in mM): 130 CsMeSO_4_, 5 CsCl, 4 NaCl, 1 MgCl_2_, 10 HEPES, 5 EGTA, 5 QX-314 Cl, 0.5 GTP, 10 Na-phosphocreatine, 5 MgATP, 0.1 Spermine and 181 units Creatine phosphokinase (290 mOsm, pH adjusted to 7.3 with CsOH). For current clamps experiments and sodium/potassium currents recordings, pipettes were filled with a potassium-based solution containing (in mM): 145 K-gluconate, 3 NaCl, 1 MgCl_2_, 1 EGTA, 0.3 CaCl_2_, 2 Na-ATP, 0.3 Na-GTP, 0.2 cAMP and 10 HEPES (290 mOsm, pH adjusted to 7.3 with KOH). Patch pipettes displayed a resistance of 4–7 MΩ.

### *Synaptic events recordings*: mEPSCs and mIPSCs were recorded in voltage clamp mode, holding cells at -70 and 10mV respectively. In the external solution 1µM of Tetrodotoxin (TTX, Tocris) and 100 µM picrotoxin (PTX, Tocris) were added for mEPSCs recordings and 1µM of Tetrodotoxin (TTX) and 50 µM of 6-cyano-7-nitroquinoxaline-2,3-dione (CNQX, Tocris) for mIPSCs recordings. Synaptic events were sampled for 5min and then exported to Clampfit 10.7 software with which the amplitude and frequency of synaptic events were quantified (threshold 6pA); all events were checked by eye and monophasic events were used for amplitude and decay kinetics, while others were suppressed but included in frequency counts as in these studies(4, 5). Tolerance for series resistance (Rs) was <35 MΩ and uncompensated; ΔRs tolerance cut-off was <20%.

### *Sodium and potassium currents*: Sodium and potassium currents were acquired in voltage-clamp mode. Sodium channel currents are reported as inward peak currents and potassium channel currents as outward currents during series of voltage steps of 10mV from -70mV to 20mV.

### *Neuronal excitability*: for assessing neuronal excitability, action potential (AP) firing was recorded in current-clamp mode in response to incremental, depolarizing current injections of 1s duration (20pA increment of 15 steps). The number of AP firing was plotted to the corresponding current steps using Clampfit 10.7 software. In current-clamp mode, the resting membrane potential of all cells was adjusted to -65mV by injection of a small negative current if needed.

**Ratiometric measurement of calcium transients in cortical neurons**

Mouse cortical neurons (17-20DIV) treated or not with lithium (1.5mM) for 7 days were loaded with 5μM Fura-2 AM (Molecular Probes, Life Technologies) in 0.1 % BSA for 40 min, then washed for 30 min with extracellular solution containing in mM: 152 NaCl, 5 KCl, 2 CaCl_2_, 1 MgCl_2_, 10 HEPES and 10 glucose, pH 7.4, with or without LiCl at 37°C in 5% CO_2_. Regions of interest (fluorescent neurons) were selected using a Nikon TE2000-U inverted microscope. Fura-2 was excited at 340 nm and 380 nm every second and emission at 510 nm was detected with a high-resolution cooled CCD camera (Cool Snap HQ, Roper Scientific-Photometrics) interfaced to a Pentium III PC. Changes in intracellular calcium levels were determined ratiometrically (∆ 340/380 ratios) using the MetaFluor 7.0 software (Molecular Devices). For each dish, agonist-induced increase in intracellular calcium was measured by subtracting the baseline ratio (F_0_) from the peak ratio (F) of the response (∆F = F-F_0_). During recording, cells were constantly perfused with extracellular solution and subjected to different experimental conditions. All recordings were conducted at room temperature and cells were constantly exposed to TTX (1μM) and picrotoxin (100μM) or TTX + picrotoxin + glycine (1μM, Sigma), depending on the experimental conditions. L-glutamic acid (1μM, Sigma) was used as agonist for metabotropic and ionotropic receptors activation. NMDA (10μM, Sigma) was used for selective NMDA receptor-channel activation. DHPG (100μM, Sigma), in the presence of the NMDA antagonist amino-5-phosphonovaleric acid (AP-5, 50μM, Sigma), was used to activate mGluR1 and mGluR5 receptors.

### RNA extraction and RNA sequencing

Two cortical neuronal cultures chronically treated or not with LiCl (1 x10^6^ cells for each sample) were collected at 18 DIV. Total RNA was extracted using miRNeasy kit (Qiagen, USA) according the manufacturer’s instructions. RNA was resuspended in RNAse-free water. The RNA concentration was measured on the Synergy H4 microplate reader. RNA was sent to Macrogen Inc. for sequencing. Library preparation was done using the TruSeq Stranded Total RNA Kit (Ilumina) with Ribo-Zero depletion. Sequencing was done on the NovaSeq 6000 at 150bp paired end reads with a total of 113M reads.

### Differential expression analysis and Pathway enrichment

In brief, Salmon was used to pseudo-align FASTQ files against Ensembl v94(6). A likelihood ratio test was used to identify differentially expressed genes with sleuth, and a Wald test was used to get a beta-estimate(7). P-values were corrected using for false-discovery rate (FDR) via Benjamini-Hochberg procedure. Enrichment in pathways and gene sets were investigated using Gene Network (genenetwork.nl)(8). For a detailed methodology on RNAseq processing, please see Liao et al. (2019)(9).

**Supplementary Figures**

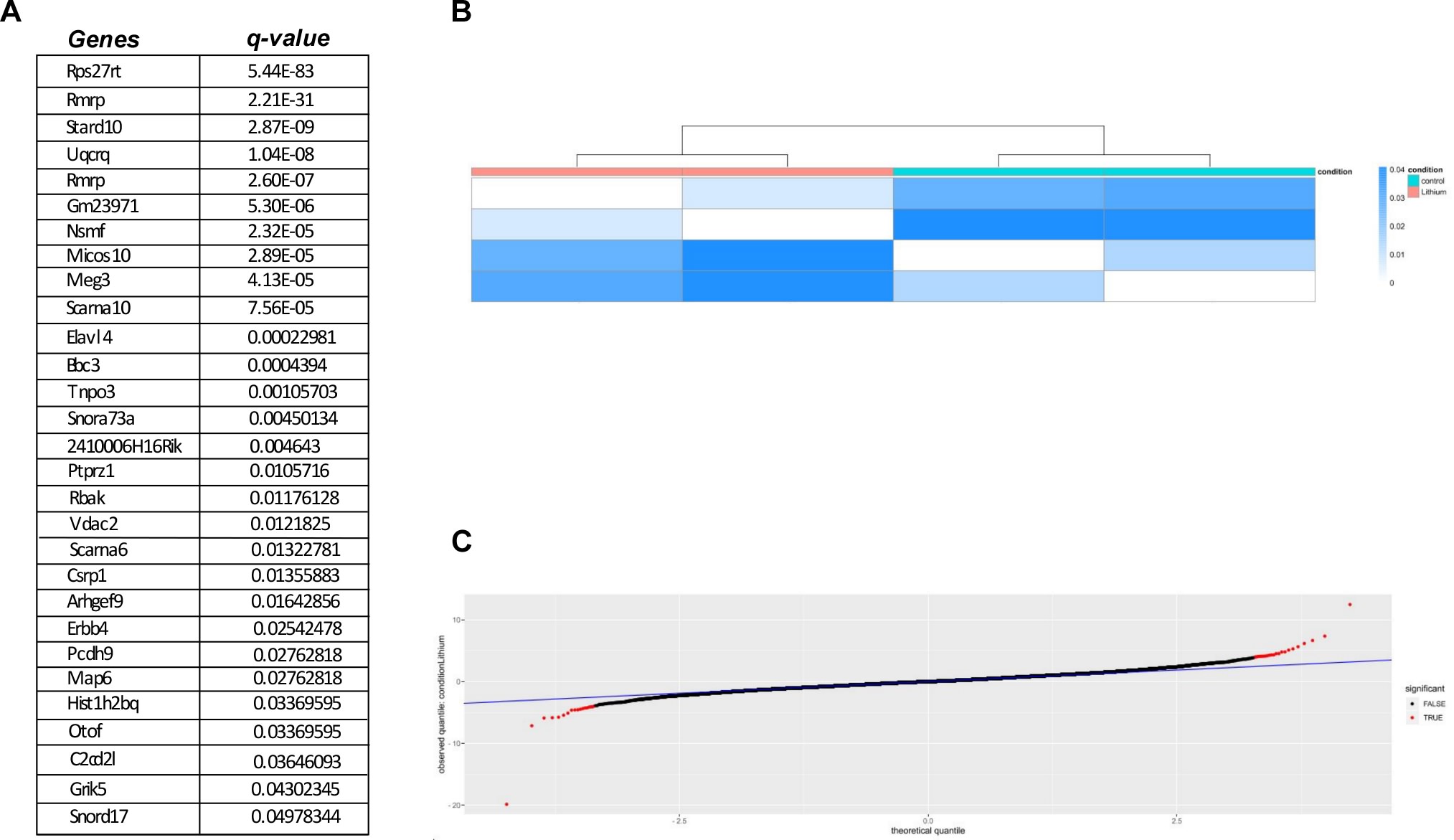

**Supplementary figure 1: *Differentially expressed genes in lithium treated vs control cortical mouse*** neurons. **A)** List of genes that are differentially expressed in neurons chronically treated with LiCl (1.5mM) for 7 days. **B)** Heat map of lithium and control samples. **C)** QQ-plot of RNAseq differential expression data. Blue line shows expected values.

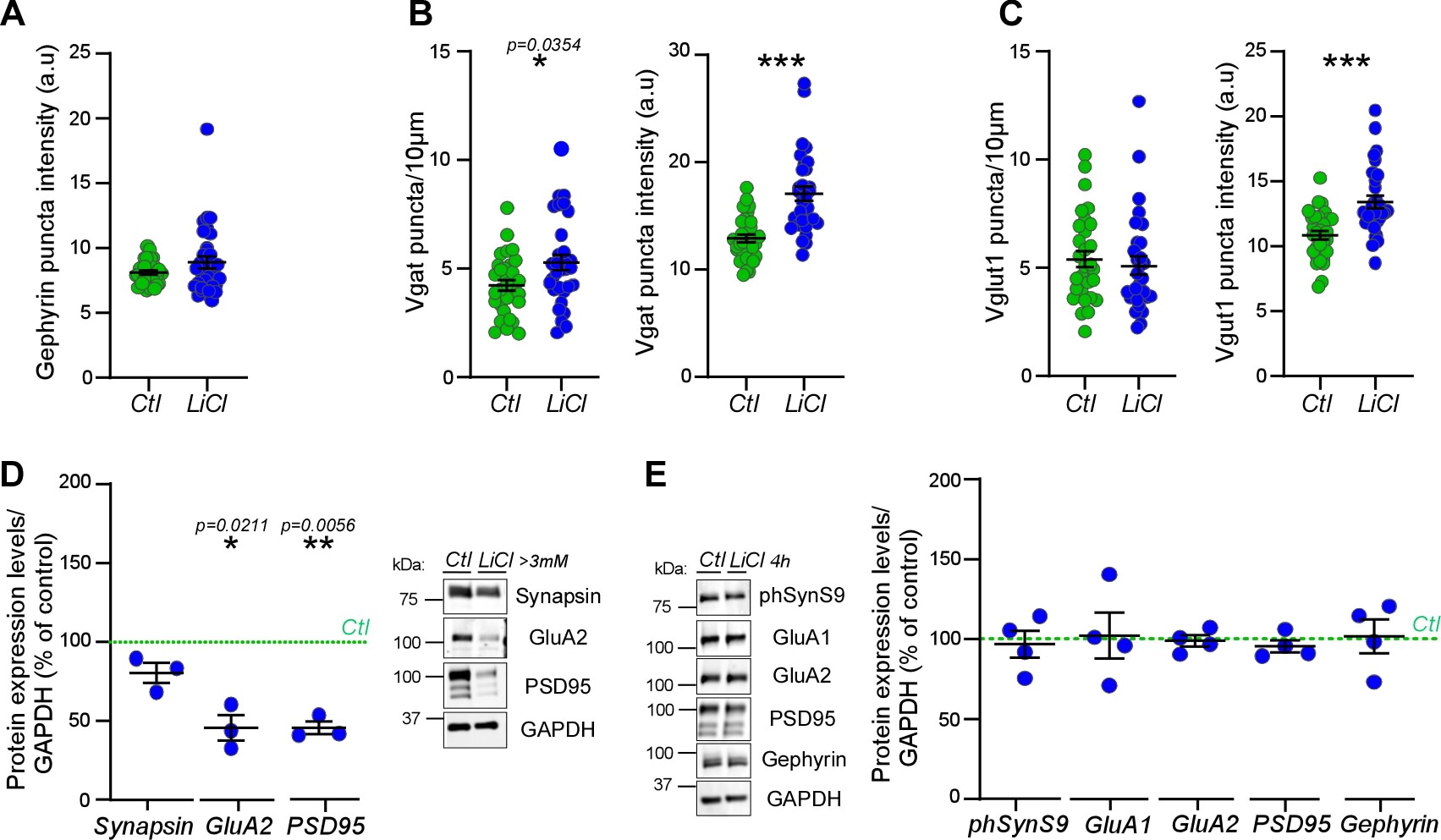

**Supplementary figure 2: *Excitatory and inhibitory synaptic changes by chronic lithium treatment.***

**A)** Scatter plot show quantification of Gephyrin puncta intensity in neurons treated or not with LiCl. **B)** Scatter plot show quantification of Vgat puncta density/10µm and Vgat puncta intensity of secondary/tertiary dendrites from neurons treated or not with LiCl (1.5mM) for 7 days. N = 32 neurons per condition from three separate experiments. **c** Scatter plot show quantification of Vglut1 puncta density/10µm and Vglut1 puncta intensity of secondary/tertiary dendrites from neurons treated or not with LiCl (1.5mM) for 7 days. N = 30 neurons per condition from three separate experiments. Data shown in **A**-**C** are the mean ± s.e.m. and statistical significance was determined using non-parametric Mann-Whitney test. ***p < 0.0005

**D)** Representative immunoblot anti-Synapsin1, GluA2, PSD95 and GAPDH with scatter plot show quantification of these protein expression levels normalized with GAPDH and represented as percentage of control of 18 DIV neuronal extract from neurons treated chronically or not with LiCl (~3.5mM) from three separate experiments. **E)** Representative and quantification of some presynaptic and postsynaptic protein expression levels normalized with GAPDH and represented as percentage of control of 18 DIV neuronal extract from neurons treated or not with LiCl (1.5mM) for 4h from four separate experiments. Data shown in **D-E** are the mean ± s.e.m. and statistical significance was determined with one sample t-test with hypothetical value 100 for controls.

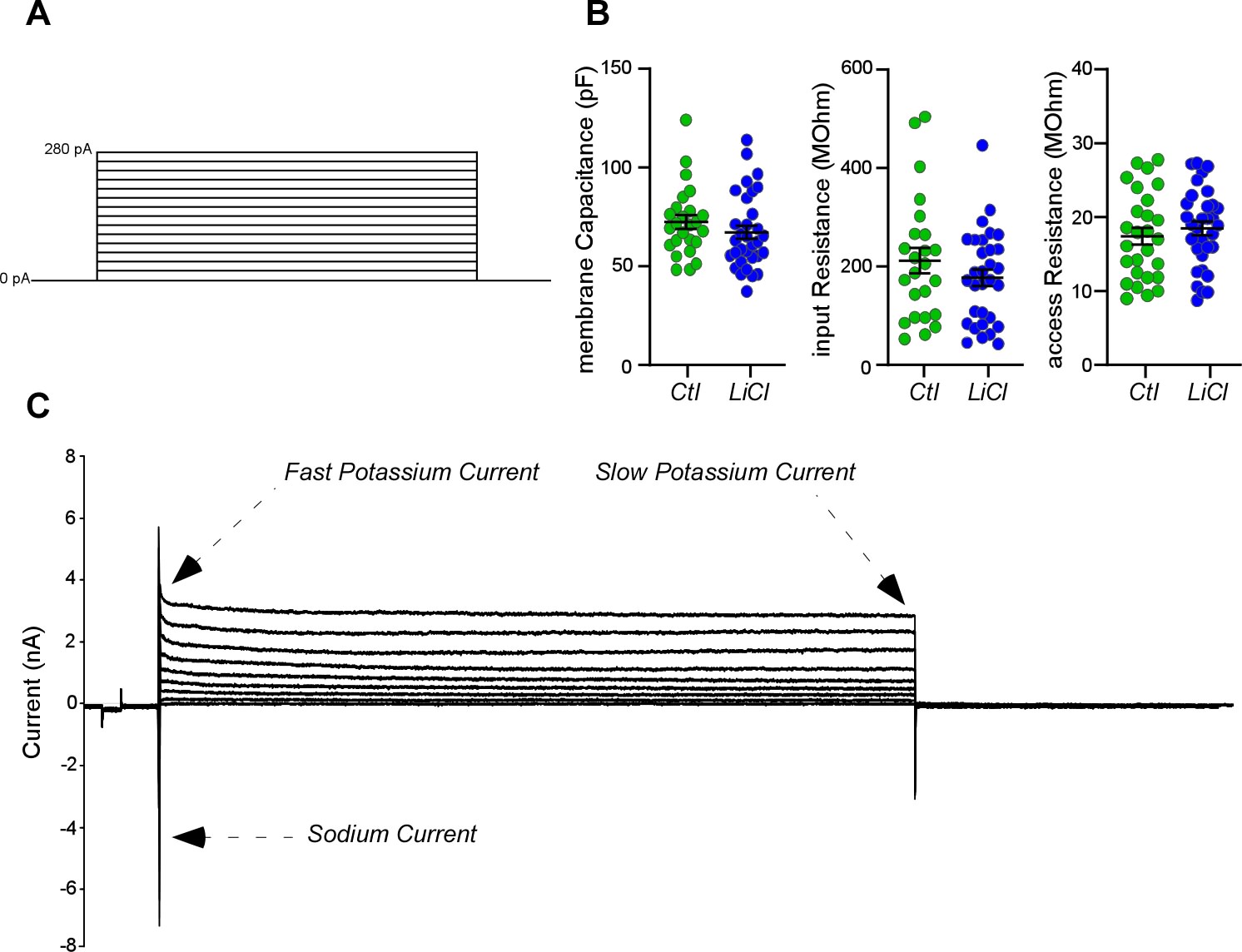

**Supplementary figure 3: *Measurements of voltage-dependent sodium and potassium currents***

**A)** Waveform stimulus protocol of 20pA increment of 14 sweeps. **B)** Scatter plots show quantification of membrane properties prior current injection steps. Data are the mean ± s.e.m. from ~32 neurons per condition from four independent experiments and statistical non-significance was determined by parametric unpaired t-test. **C)** Electrophysiological sample trace shows voltage-dependent sodium and potassium currents. Arrowheads in the positive current indicate the peak amplitude of fast and slow potassium currents. Arrowhead in the negative current indicates sodium current.

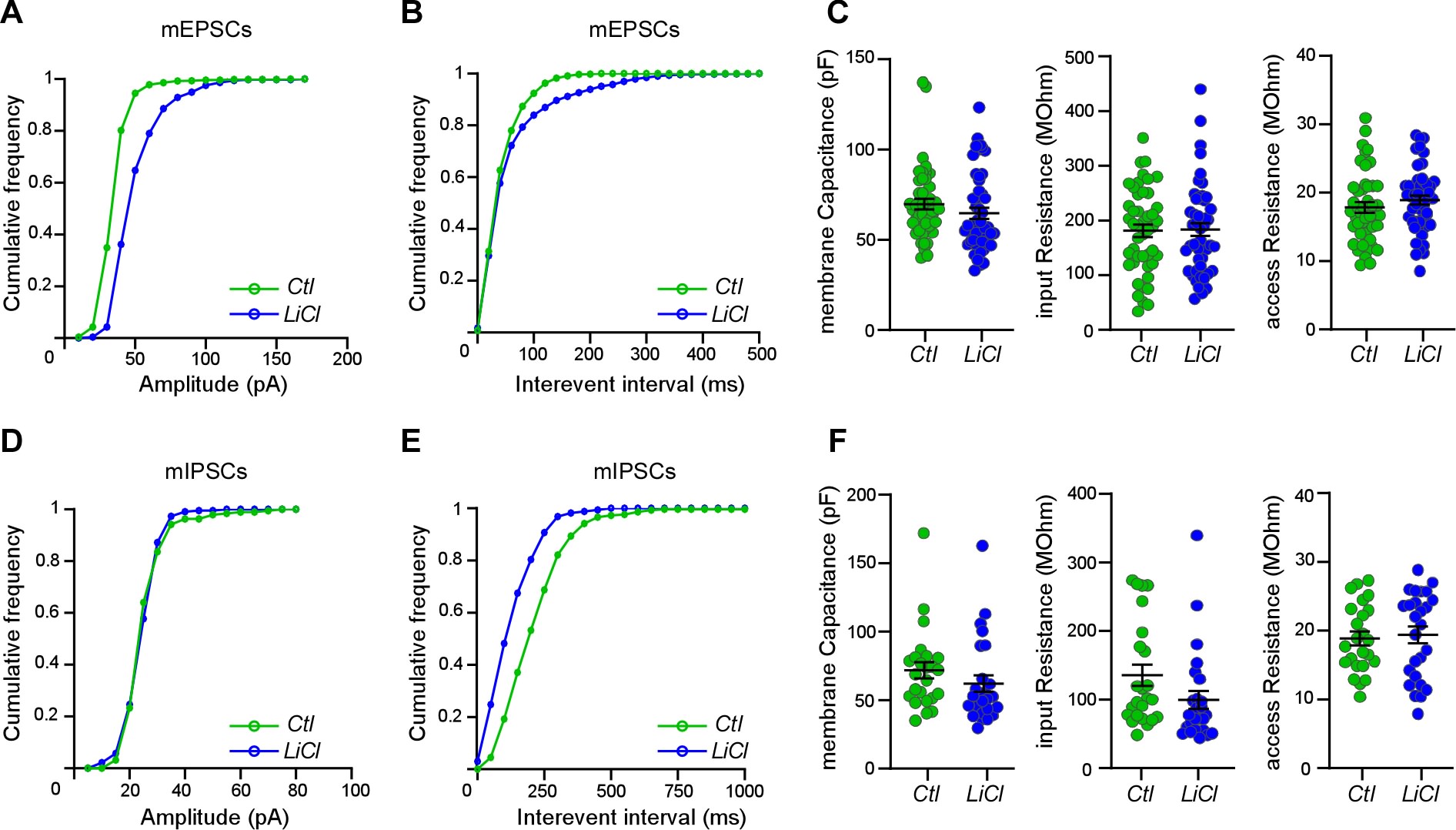

**Supplementary figure 4: *Chronic lithium treatment alters the spontaneous excitatory and inhibitory synaptic transmission.***

Cumulative frequency for amplitude (**A** and **D)** and interevent intervals **(B** and **E)** of mEPSCs and mIPSCs respectively recorded from neurons treated or not chronically with LiCl (1.5mM). ~45 neurons from four independent experiments for mEPSCs and ~26 neurons from three independent experiments for mIPSCs. **C** and **F** Scatter plots show quantification of membrane properties Data are the mean ± s.e.m. and statistical non-significance was determined by parametric unpaired t-test.

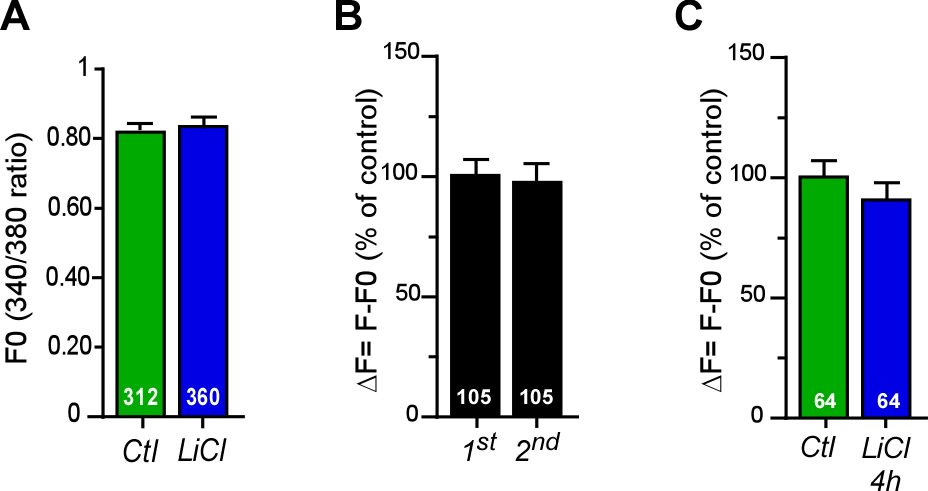

**Supplementary figure 5:** ***Acute lithium treatment does not affect glutamate-induced calcium flux.***

**A)** Averaged baseline calcium levels prior drugs stimulations. **B)** Quantification of calcium changes as percentage of control upon two repeated glutamate (1µM) exposures in control neurons. **C)** Quantification of calcium changes as percentage of control upon glutamate (1µM) stimulation in mouse primary cortical neurons treated or not with LiCl (1.5mM) for 4h. Number of neurons is indicated on each histogram from 3 independent experiments. Data shown in **A-C** are the mean ± s.e.m. and statistically non-significant with parametric unpaired t-tests.

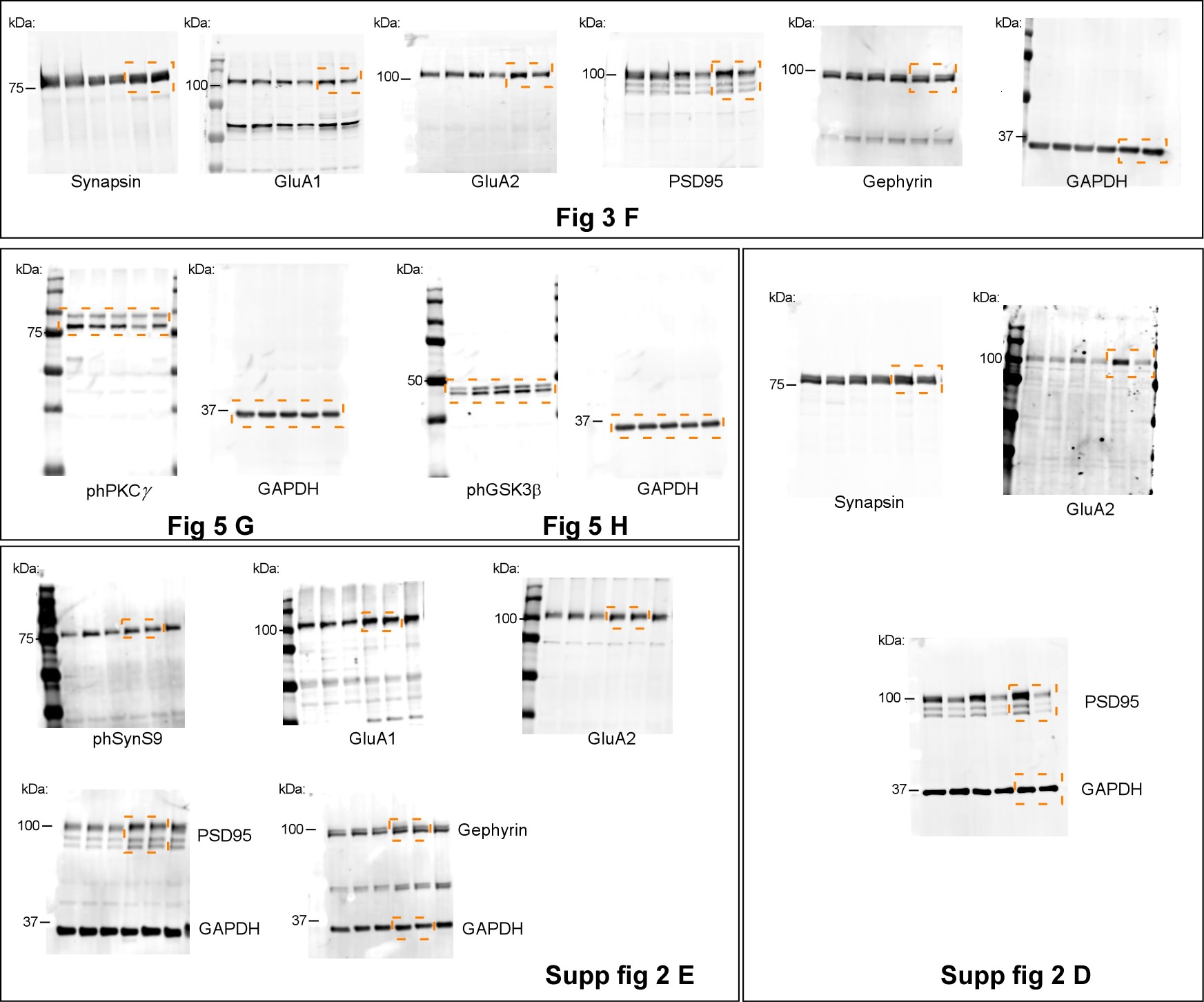

**Supplementary figure 6: *Original uncropped blots***. Orange boxed regions represent the portion used in the indicated figures.
